## Supplementary Information for "Effect of alpha-tubulin acetylation on the doublet microtubule structure"

### Supplementary Figures and Tables

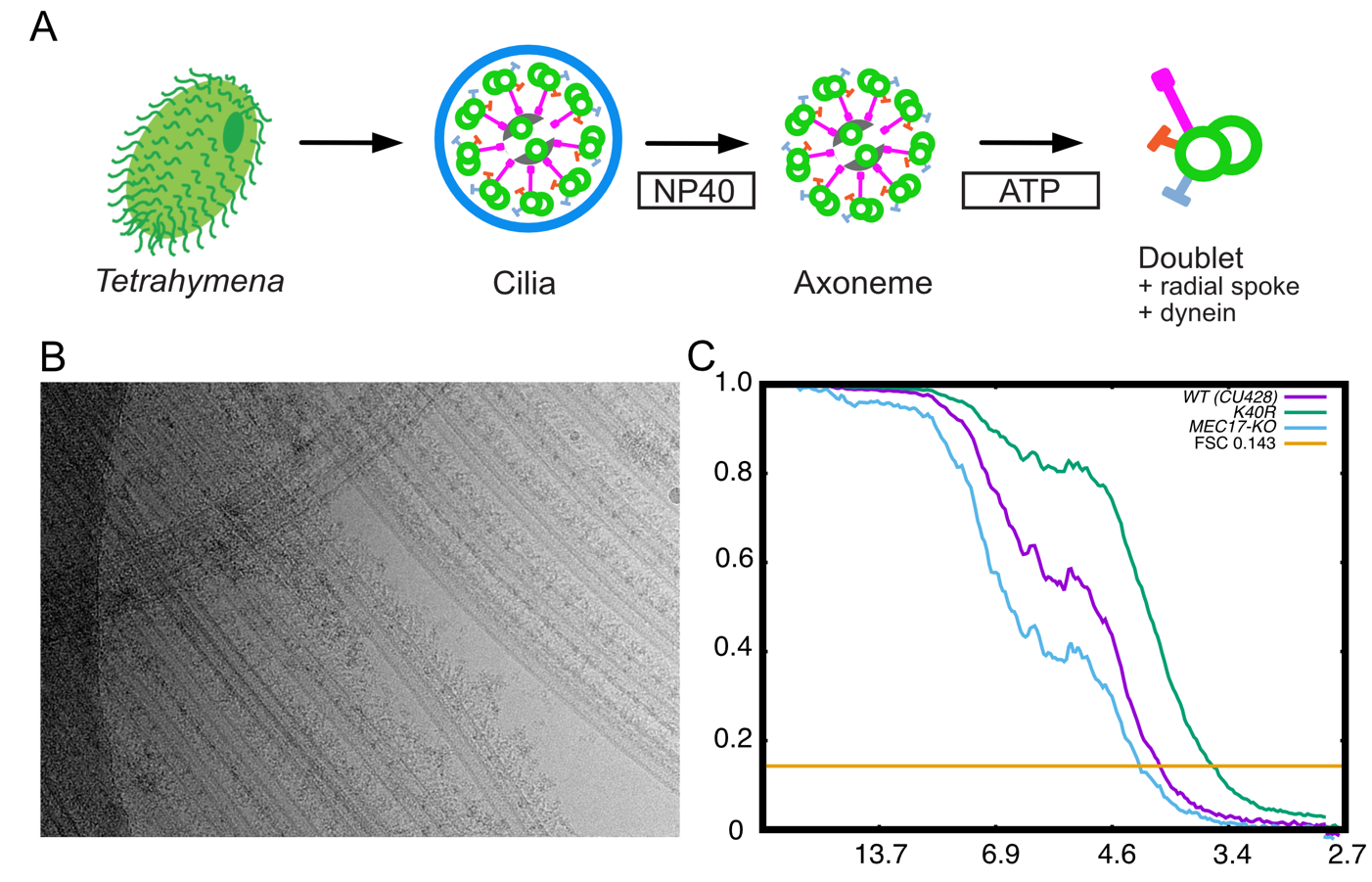

**Supplementary Fig. 1. Purification and structure determination of DMT.**

(A) Purification scheme of the axoneme in this study. (B) A typical cryo-EM image of the *Tetrahymena* DMT. (C) Gold-standard Fourier shell correlation of the 48-nm repeat cryo-EM maps of the *WT, K40R, and MEC17-KO* *Tetrahymena* strains.

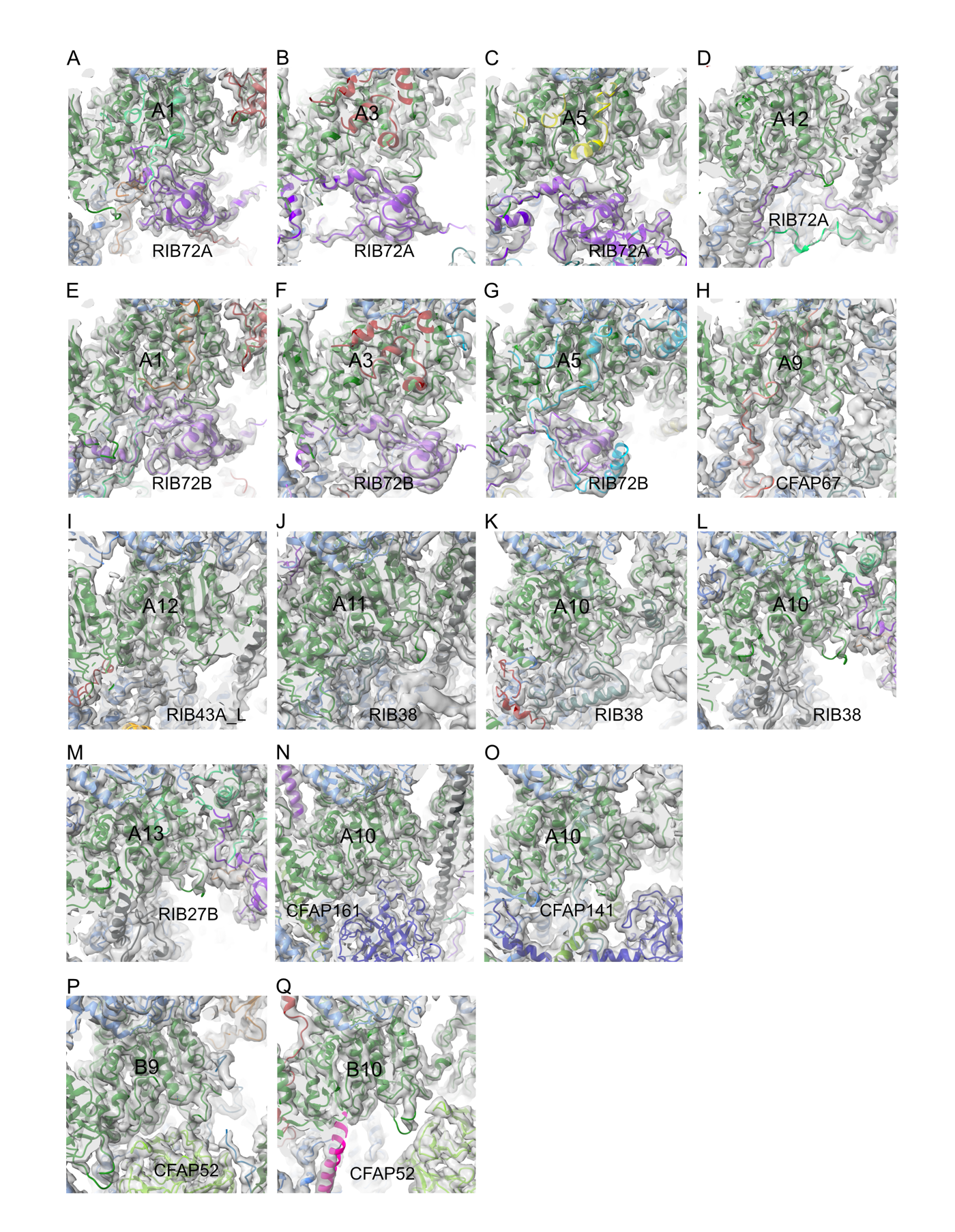
**Supplementary Fig. 2. Diverse structures of the αK40 loops.** (A-Q) Cryo-EM maps and models of partial and full αK40 loops interacting with different MIPs.

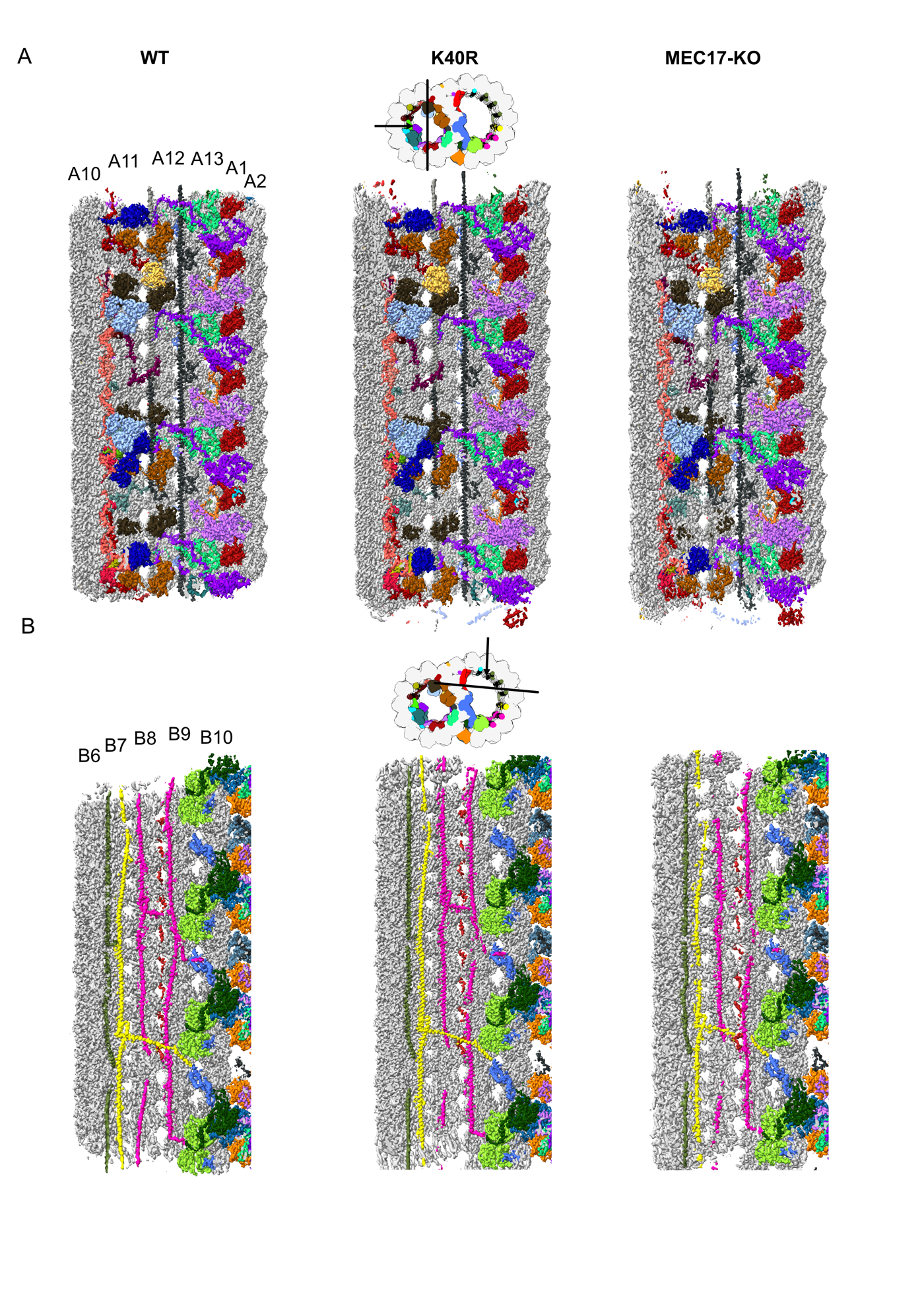

**Supplementary Fig. 3**. **Decoration of MIPs are similar in 48-nm cryo-EM maps of the DMT from WT, K40R and MEC17-KO.** (A) View from inside the A-tubule. (B) View from inside B-tubule.

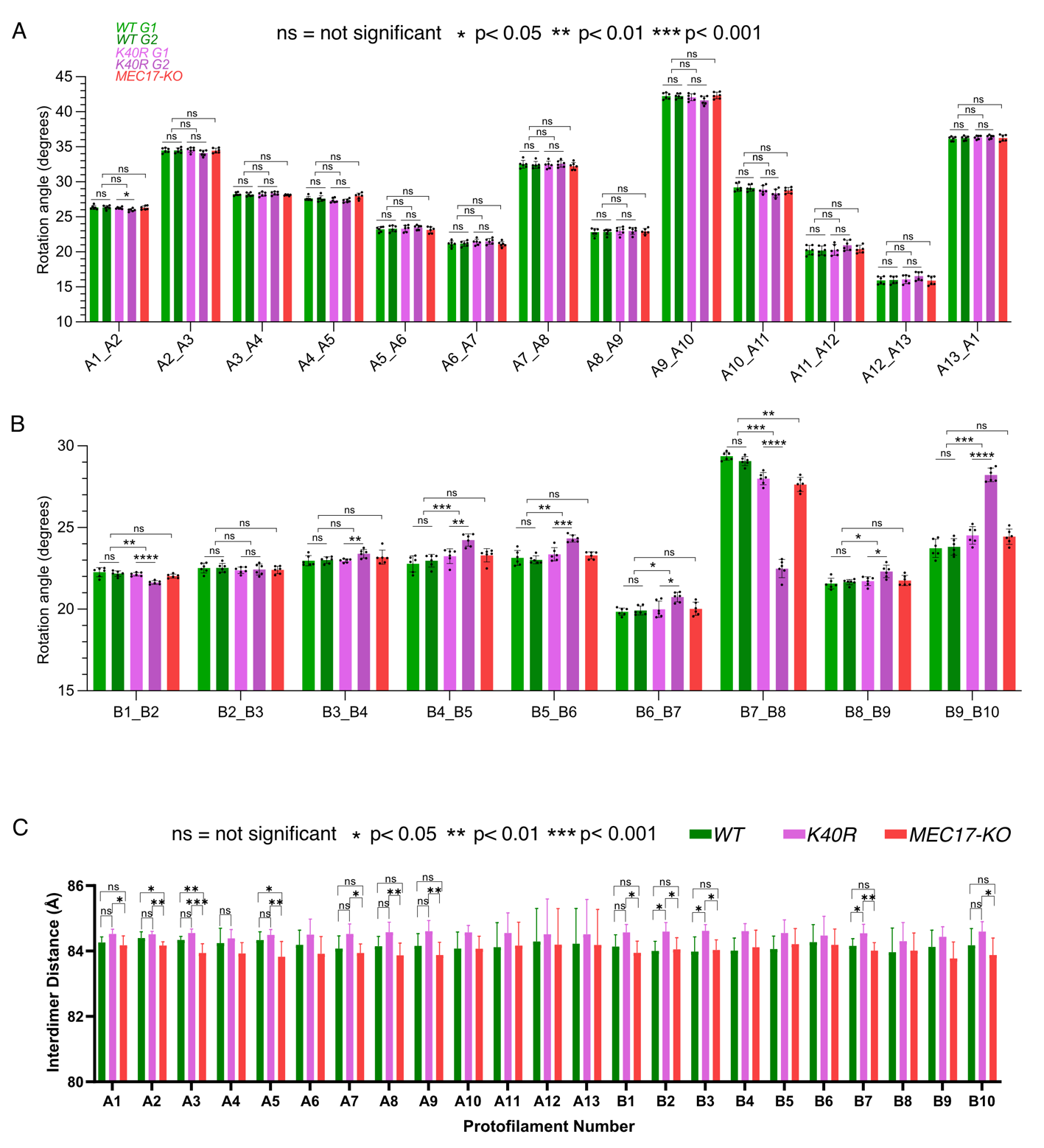

**Supplementary Fig. 4. Tubulin lattice measurement from the DMT from *WT, K40R*, and *MEC17-KO* *Tetrahymena* species.** (A) Inter-PF rotation angle between A-tubule protofilament pairs. (B) Inter-PF rotation angle between B-tubule protofilament pairs. For (A) and (B), normality was assessed with a Shapiro-Wilk test. Unpaired t tests or Mann Whitney tests were carried out for each angle between both WT and K40R groups separately. One-way ANOVA or Kruskal Wallis tests were carried out for each angle between combined WT points, combined K40R points and MEC17-KO. ns: not significant, * p-value < 0.05, **<0.01, *** <0.001 and **** <0.0001. (C) Interdimer distance within the same PF measured from cryo-EM density maps of *WT, K40R*, and *MEC17-KO* *Tetrahymena* species.

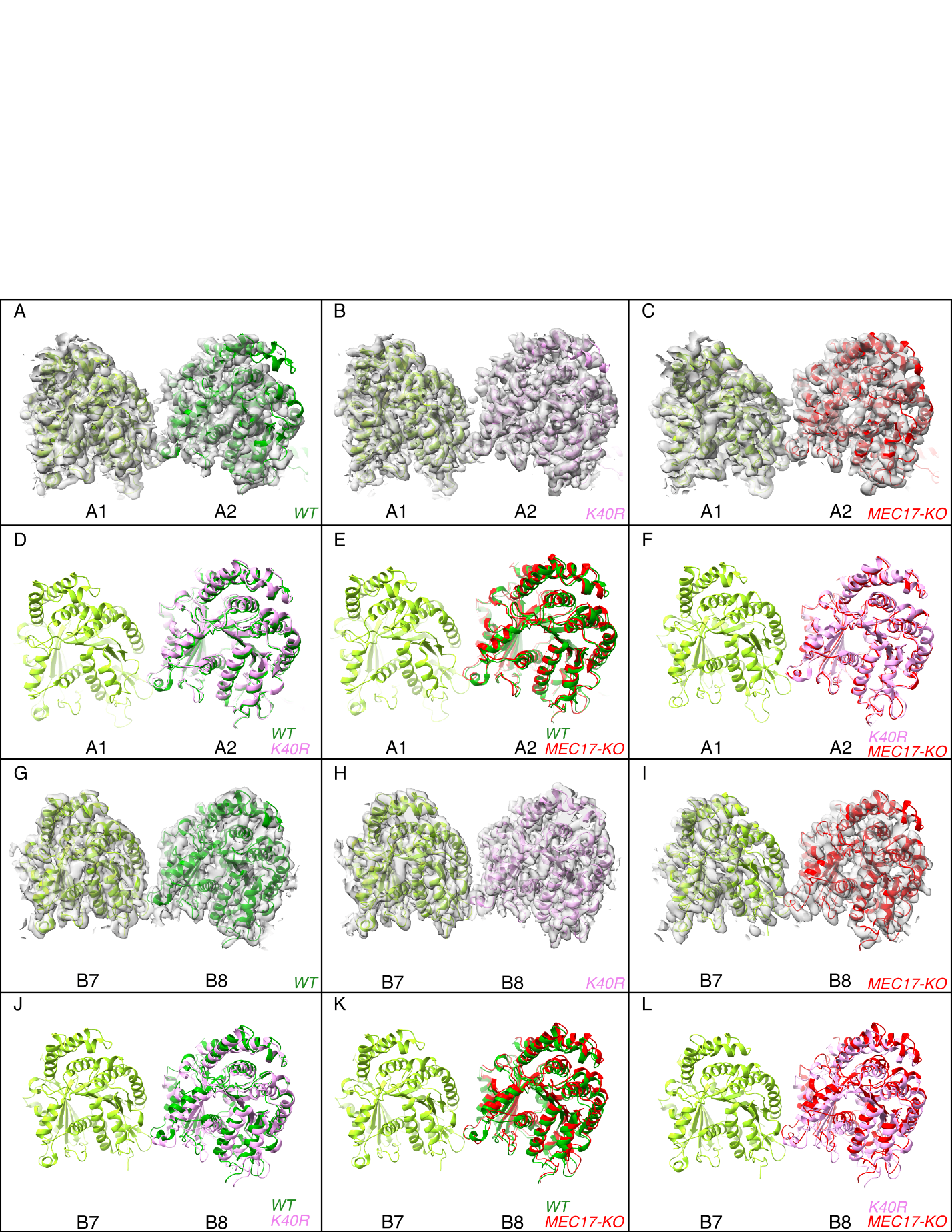

**Supplementary Fig. 5: Deacetylation affects the inter-PF angles in the DMT.**

(A-F) Comparison of inter-PF rotation angle change between A1 and A2, showing statistically non-significant changes. (G-L) Comparison of inter-PF rotation angle change between B7 and B8, showing significant changes.

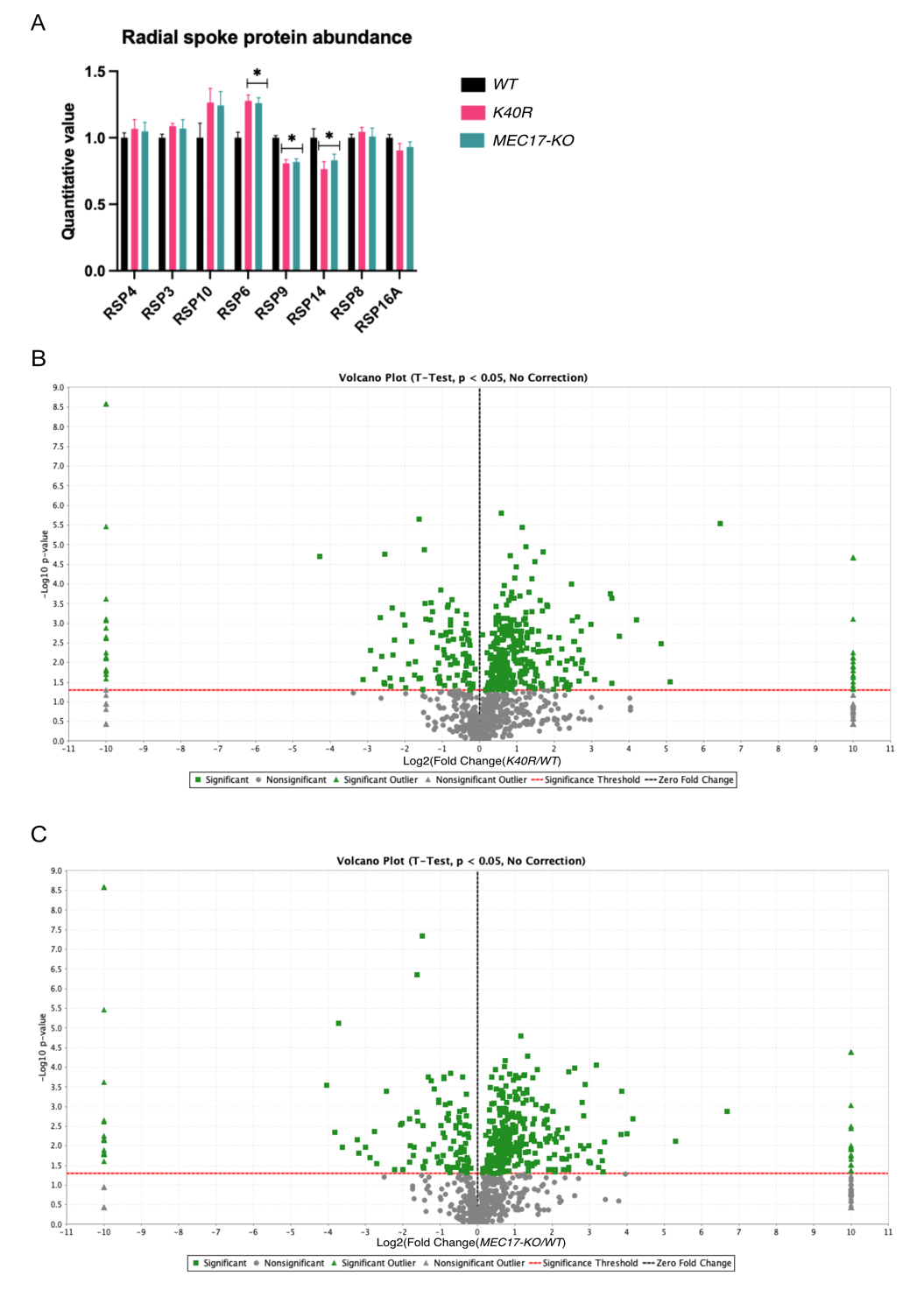

**Supplementary Fig. 6. Mass spectrometry analysis of *WT*, *K40R* and *MEC17-KO***

(A) Radial spoke protein abundance in mass spectrometry triplicates of *WT*, *K40R*, and *MEC17-KO* *Tetrahymena* species. Asterisk (*) indicates a significant difference with p < 0.05. (B) Volcano plot showing up- and downregulated proteins in *K40R* mutants compared against *WT*, y-axis showing the -Log10 p values, and x-axis showing the Log2-fold change where negative and positive values suggest potential downregulation and upregulation, respectively. (C) Volcano plot showing up- and downregulated proteins in *MEC17-KO* mutants compared to *WT*.

**Supplementary Table 1.** Count of full, partial, and missing αK40 loops based upon the *K40R* cryo-EM density maps, along with possible MIP interactions for αK40 loops in each PF.

| **PF(48-nm Internal Repeat)** | **Full** α**K40 Loop Count** | **Partial** α**K40 Loop Count** | **Possible MIP Interaction** |
| --- | --- | --- | --- |
| A1 | 6 | 0 | RIB72A, RIB72B |
| A2 | 0 | 0 | RIB72A, RIB72B, CFAP115 |
| A3 | 6 | 0 | RIB72A, RIB72B |
| A4 | 0 | 3 | RIB72A, RIB72B |
| A5 | 6 | 0 | RIB72A, RIB72B |
| A6 | 1 | 1 | STPG2 |
| A7 | 1 | 0 | CFAP127 |
| A8 | 0 | 0 |  |
| A9 | 3 | 0 | CFAP67, CFAP161A, CFAP129, RIB57 |
| A10 | 4 | 1 | CFAP141, CFAP143, CFAP161A, RIB38 |
| A11 | 3 | 2 | CFAP143, CFAP161, RIB38, RIB57 |
| A12 | 5 | 1 | RIB43A_L, RIB72A |
| A13 | 1 | 3 | RIB72B, RIB27B |
| B1 | 3 | 0 | CFAP77, OJ2 |
| B2 | 0 | 0 |  |
| B3 | 0 | 0 |  |
| B4 | 0 | 0 |  |
| B5 | 0 | 0 |  |
| B6 | 0 | 0 |  |
| B7 | 0 | 0 |  |
| B8 | 0 | 0 |  |
| B9 | 3 | 0 | CFAP52, CFAP210, CFAP45 |
| B10 | 3 | 0 | CFAP52, PACRGA |

**Supplementary Table 2.** Inter-PF angles between subsequent PF obtained from cryo-EM maps of *WT*, *K40R*, and *MEC17-KO* *Tetrahymena* cilia.

|  | ***WT*** | | ***K40R*** | | ***MEC17-KO*** | |
| --- | --- | --- | --- | --- | --- | --- |
| **PF pair** | **Mean rotation angle (degrees)**  **n = 6** | **Standard Deviation** | **Mean rotation angle (degrees)**  **n = 6** | **Standard Deviation** | **Mean rotation angle (degrees)**  **n = 6** | **Standard Deviation** |
| A1_A2 | 26.454 | 0.230 | 25.987 | 0.415 | 26.356 | 0.281 |
| A2_A3 | 34.578 | 0.318 | 34.270 | 0.338 | 34.391 | 0.364 |
| A3_A4 | 28.017 | 0.238 | 28.247 | 0.262 | 28.279 | 0.475 |
| A4_A5 | 27.823 | 0.522 | 27.716 | 0.422 | 28.152 | 0.615 |
| A5_A6 | 22.892 | 0.633 | 23.306 | 0.416 | 22.996 | 0.546 |
| A6_A7 | 21.501 | 0.671 | 21.392 | 0.349 | 21.138 | 0.362 |
| A7_A8 | 32.418 | 0.570 | 32.577 | 0.577 | 32.339 | 0.528 |
| A8_A9 | 22.918 | 0.535 | 22.930 | 0.413 | 22.971 | 0.399 |
| A9_A10 | 41.819 | 0.604 | 41.593 | 0.656 | 42.305 | 0.431 |
| A10_A11 | 29.368 | 0.610 | 28.645 | 0.626 | 28.851 | 0.438 |
| A11_A12 | 20.461 | 0.520 | 20.878 | 0.573 | 20.311 | 0.557 |
| A12_A13 | 15.875 | 0.491 | 16.399 | 0.581 | 16.007 | 0.626 |
| A13_A1 | 35.974 | 0.370 | 36.050 | 0.481 | 36.173 | 0.464 |
| B1_B2 | 22.161 | 0.240 | 21.737 | 0.158 | 22.059 | 0.141 |
| B2_B3 | 22.579 | 0.272 | 22.384 | 0.252 | 22.385 | 0.217 |
| B3_B4 | 22.919 | 0.193 | 23.447 | 0.283 | 23.258 | 0.439 |
| B4_B5 | 23.137 | 0.448 | 23.839 | 0.397 | 23.232 | 0.407 |
| B5_B6 | 23.211 | 0.639 | 24.329 | 0.245 | 23.359 | 0.261 |
| B6_B7 | 20.003 | 0.468 | 20.688 | 0.361 | 19.951 | 0.418 |
| B7_B8 | 29.039 | 0.351 | 23.353 | 0.624 | 27.737 | 0.481 |
| B8_B9 | 21.829 | 0.385 | 22.303 | 0.466 | 21.939 | 0.554 |
| B9_B10 | 23.554 | 0.455 | 27.409 | 0.417 | 24.556 | 0.527 |

**Supplementary Table 3**. ANOVA test p values and significance of the interdimer distances of tubulin subunits within each PF in *WT,* *K40R*, and *MEC17-KO* *Tetrahymena* cilia. ns: not significant, * p-value < 0.05, **<0.01, *** <0.001 and **** <0.0001.

| **PF** | **Interdimer distances from *WT* (Å) (mean ± SD, n = 6)** | **Interdimer distances from *K40R* (Å) (mean ± SD, n = 6)** | **Interdimer distances from *MEC17-KO* (Å) (mean ± SD, n = 6)** | **ANOVA p values** | **p value summary** |
| --- | --- | --- | --- | --- | --- |
| A1 | 84.352 ± 0.276 | 84.518 ± 0.137 | 84.194 ± 0.221 | 0.097 | ns |
| A2 | 84.453 ± 0.144 | 84.512 ± 0.083 | 84.222 ± 0.085 | 0.002 | ** |
| A3 | 84.426 ± 0.301 | 84.548 ± 0.113 | 84.218 ± 0.090 | 0.048 | * |
| A4 | 84.300 ± 0.451 | 84.390 ± 0.244 | 84.016 ± 0.210 | 0.190 | ns |
| A5 | 84.506 ± 0.268 | 84.493 ± 0.151 | 84.080 ± 0.195 | 0.009 | ** |
| A6 | 84.479 ± 0.473 | 84.495 ± 0.441 | 84.105 ± 0.388 | 0.305 | ns |
| A7 | 84.418 ± 0.537 | 84.519 ± 0.282 | 84.091 ± 0.246 | 0.207 | ns |
| A8 | 84.367 ± 0.298 | 84.575 ± 0.280 | 84.093 ± 0.191 | 0.034 | * |
| A9 | 84.393 ± 0.428 | 84.603 ± 0.301 | 84.132 ± 0.351 | 0.157 | ns |
| A10 | 84.364 ± 0.331 | 84.567 ± 0.195 | 84.141 ± 0.286 | 0.082 | ns |
| A11 | 84.352 ± 0.639 | 84.546 ± 0.565 | 84.166 ± 0.636 | 0.629 | ns |
| A12 | 84.323 ± 1.063 | 84.511 ± 0.989 | 84.198 ± 1.000 | 0.888 | ns |
| A13 | 84.355 ± 0.957 | 84.506 ± 0.977 | 84.204 ± 1.018 | 0.890 | ns |
| B1 | 84.253 ± 0.301 | 84.566 ± 0.226 | 84.136 ± 0.296 | 0.068 | ns |
| B2 | 84.171 ± 0.302 | 84.592 ± 0.257 | 84.106 ± 0.343 | 0.046 | * |
| B3 | 84.174 ± 0.413 | 84.617 ± 0.175 | 84.112 ± 0.255 | 0.034 | * |
| B4 | 84.134 ± 0.359 | 84.607 ± 0.211 | 84.185 ± 0.417 | 0.085 | ns |
| B5 | 84.036 ± 0.515 | 84.547 ± 0.368 | 84.133 ± 0.512 | 0.223 | ns |
| B6 | 84.419 ± 0.358 | 84.473 ± 0.538 | 84.090 ± 0.617 | 0.465 | ns |
| B7 | 84.042 ± 0.356 | 84.540 ± 0.252 | 84.271 ± 0.177 | 0.035 | * |
| B8 | 84.129 ± 0.494 | 84.299 ± 0.525 | 84.061 ± 0.516 | 0.754 | ns |
| B9 | 84.259 ± 0.371 | 84.432 ± 0.285 | 84.062 ± 0.446 | 0.321 | ns |
| B10 | 84.359 ± 0.575 | 84.590 ± 0.281 | 84.194 ± 0.507 | 0.431 | ns |

**Supplementary Table 4**. Proteins upregulated at least twofold in the *K40R* mutant compared to *WT.*

| **Gene name** | **Gene ID** | **UniProt ID** | **Avg *WT*** | **Avg *K40R*** | **p value** | **Average fold change** | **Homolog in *C. reinhardtii*** | **Homolog in *Homo sapiens*** |
| --- | --- | --- | --- | --- | --- | --- | --- | --- |
| WD domain, G-beta repeat protein | TTHERM_00006270 | Q22SA9 | 1.160 | 4.649 | 0.047 | 4.000 | NA | NA |
| Uncharacterized protein | TTHERM_00449680 | Q238V2 | 0.460 | 1.932 | 0.026 | 4.200 | NA | NA |
| Uncharacterized protein | TTHERM_00703420 | Q22GI1 | 3.250 | 13.702 | 0.035 | 4.200 | NA | NA |
| Uncharacterized protein | TTHERM_00392650 | Q233I4 | 1.860 | 8.598 | 0.010 | 4.400 | NA | NA |
| Uncharacterized protein | TTHERM_00548020 | I7MDA2 | 1.860 | 8.598 | 0.005 | 4.600 | NA | NA |
| SelT/selW/selH selenoprotein domain protein | TTHERM_00195980 | Q23K36 | 0.460 | 2.289 | 0.042 | 4.900 | NA | NA |
| FAM184 domain-containing protein | TTHERM_01321540 | Q232U4 | 0.930 | 4.649 | 0.033 | 5.000 | NA | [Q8NB25](https://www.uniprot.org/uniprot/Q8NB25) |
| Calmodulin | NA | P02598 | 0.930 | 4.734 | 0.038 | 5.100 | NA | Calmodulin 1,2,3 |
| Dynein light chain 8-like E | NA | Q1HFV9 | 0.930 | 5.006 | 0.036 | 5.400 | NA | [P63167](https://www.uniprot.org/uniprot/P63167) |
| Uncharacterized protein | TTHERM_00847020 | Q22US8 | 0.700 | 3.927 | 0.008 | 5.600 | NA | NA |
| Kinesin-like protein | TTHERM_01347910 | Q229Q4 | 0.470 | 2.700 | 0.003 | 5.800 | NA | [Q2TAC6](https://www.uniprot.org/uniprot/Q2TAC6) |
| Uncharacterized protein | TTHERM_00483660 | I7M1N4 | 0.460 | 2.700 | 0.003 | 5.800 | NA | NA |
| Cyclic nucleotide-binding domain protein | TTHERM_00841230 | I7M4G5 | 0.460 | 2.700 | 0.003 | 5.800 | NA | NA |
| Kinesin-like protein | TTHERM_00758880 | Q23JL7 | 0.460 | 2.747 | 0.011 | 5.900 | [A0A2K3D1C7](https://www.uniprot.org/uniprot/A0A2K3D1C7) | [A0A7I2V3N5](https://www.uniprot.org/uniprot/A0A7I2V3N5) |
| Scp-like extracellular protein, | TTHERM_00145900 | I7MI85 | 1.620 | 10.088 | 0.001 | 6.200 | NA | NA |
| Uncharacterized protein | TTHERM_00659030 | I7M3P7 | 1.160 | 7.762 | 0.014 | 6.700 | NA | NA |
| Hect domain and RCC1-like domain protein | TTHERM_00535910 | I7LX15 | 0.700 | 4.696 | 0.010 | 6.800 | NA | NA |
| Uncharacterized protein | TTHERM_00827160 | Q22EF3 | 0.460 | 3.159 | 0.009 | 6.800 | [A0A2K3DZJ7](https://www.uniprot.org/uniprot/A0A2K3DZJ7) | NA |
| Uncharacterized protein | TTHERM_000732839 | W7XGQ6 | 0.930 | 7.341 | 0.001 | 7.900 | NA | NA |
| Uncharacterized protein | TTHERM_00939090 | Q22DN4 | 0.230 | 1.987 | 0.027 | 8.500 | NA | NA |
| Uncharacterized protein | TTHERM_00388510 | Q23RD8 | 1.160 | 13.239 | 0.000 | 11.000 | NA | NA |
| B-box zinc finger protein | TTHERM_00112790 | Q22Z92 | 0.230 | 2.692 | 0.033 | 12.000 | NA | NA |
| Cyclic nucleotide-binding domain protein | TTHERM_00143540 | I7M0S4 | 0.230 | 3.104 | 0.002 | 13.000 | NA | NA |
| Uncharacterized protein | TTHERM_00497130 | I7M4M1 | 0.230 | 4.284 | 0.000 | 18.000 | NA | NA |
| EF hand protein | TTHERM_00833830 | Q23A35 | 0.230 | 8.017 | 0.031 | 34.000 | NA | NA |

**Supplementary Table 5.** Proteins upregulated least twofold in the *MEC17-KO* mutant compared to the *WT.*

| **Gene name** | **Gene ID** | **UniProt ID** | **Avg *WT*** | **Avg K40R** | **p value** | **Average fold change** | **Homolog in *C. reinhardtii*** | **Homolog in *Homo sapiens*** |
| --- | --- | --- | --- | --- | --- | --- | --- | --- |
| Uncharacterized protein | TTHERM_00852750 | Q24E65 | 4.180 | 16.987 | 0.001 | 4.100 | NA | NA |
| Uncharacterized protein | TTHERM_00483660 | I7M1N4 | 0.460 | 1.883 | 0.041 | 4.100 | NA | NA |
| Uncharacterized protein | TTHERM_00219140 | I7M2G4 | 1.620 | 6.778 | 0.007 | 4.200 | NA | NA |
| Hect domain and RCC1-like domain protein | TTHERM_00535910 | I7LX15 | 0.700 | 2.942 | 0.041 | 4.200 | NA | NA |
| Uncharacterized protein | TTHERM_00703420 | Q22GI1 | 3.250 | 13.940 | 0.008 | 4.300 | NA | NA |
| Uncharacterized protein | TTHERM_00190760 | I7M812 | 1.630 | 7.137 | 0.005 | 4.400 | NA | NA |
| Dynein light chain 8-like E | NA | Q1HFV9 | 0.930 | 4.474 | 0.006 | 4.800 | NA | P63167 |
| Uncharacterized protein | TTHERM_00085240 | Q236S4 | 0.230 | 1.121 | 0.021 | 4.800 | NA | NA |
| Uncharacterized protein | TTHERM_00670810 | I7LXT5 | 0.470 | 2.242 | 0.021 | 4.800 | NA | NA |
| Uncharacterized protein | TTHERM_00468010 | I7MAL9 | 0.930 | 4.536 | 0.022 | 4.900 | NA | NA |
| Uncharacterized protein | TTHERM_00010960 | Q22S36 | 0.690 | 3.415 | 0.040 | 4.900 | NA | NA |
| Uncharacterized protein | TTHERM_00548020 | I7MDA2 | 1.860 | 10.088 | 0.000 | 5.400 | NA | NA |
| Kinesin-like protein | TTHERM_00794640 | Q23VW7 | 0.470 | 2.592 | 0.018 | 5.500 | NA | A0A7I2V3V3 |
| Uncharacterized protein | TTHERM_00334380 | I7M866 | 0.460 | 2.539 | 0.040 | 5.500 | NA | NA |
| Cyclic nucleotide-binding domain protein | TTHERM_00841230 | I7M4G5 | 0.460 | 2.539 | 0.040 | 5.500 | NA | NA |
| Uncharacterized protein | TTHERM_00392650 | Q233I4 | 0.700 | 4.772 | 0.010 | 6.800 | NA | NA |
| Uncharacterized protein | TTHERM_00388510 | Q23RD8 | 1.160 | 8.555 | 0.000 | 7.400 | NA | NA |
| Uncharacterized protein | TTHERM_00827160 | Q22EF3 | 0.460 | 3.354 | 0.012 | 7.200 | NA | NA |
| Uncharacterized protein | TTHERM_000732839 | W7XGQ6 | 0.930 | 7.329 | 0.010 | 7.900 | NA | NA |
| Uncharacterized protein | TTHERM_00732900 | Q245C1 | 0.460 | 3.669 | 0.024 | 7.900 | NA | NA |
| Uncharacterized protein | TTHERM_001092361 | W7X1T3 | 0.230 | 1.892 | 0.028 | 8.100 | A0A2K3DJC4 | NA |
| Uncharacterized protein | TTHERM_00825250 | I7MCE2 | 0.230 | 1.892 | 0.028 | 8.100 | NA | NA |
| Uncharacterized protein | TTHERM_00194430 | Q23K92 | 0.930 | 8.555 | 0.000 | 9.200 | NA | NA |
| Uncharacterized protein | TTHERM_01028860 | Q23EF4 | 0.230 | 2.189 | 0.035 | 9.500 | NA | NA |
| Uncharacterized protein | TTHERM_00078910 | Q23FX3 | 0.470 | 4.536 | 0.014 | 9.700 | NA | NA |
| Uncharacterized protein | TTHERM_00449080 | Q239A7 | 0.470 | 4.728 | 0.024 | 10.000 | NA | NA |
| Uncharacterized protein | TTHERM_01009900 | Q23LQ6 | 0.230 | 3.363 | 0.000 | 15.000 | NA | NA |
| Protein kinase | TTHERM_00497440 | I7MB56 | 0.230 | 3.774 | 0.005 | 16.000 | NA | NA |
| Cyclic nucleotide-binding domain protein | TTHERM_00143540 | I7M0S4 | 0.230 | 4.124 | 0.002 | 18.000 | NA | NA |

**Supplementary Table 6.** Proteins downregulated at least twofold in the *K40R* mutant compared to the *WT*.

| **Gene name** | **Gene ID** | **UniProt ID** | **Average *WT*** | **Average K40R** | **p value** | **Average fold change** | **Homolog in *C. reinhardtii*** | **Homolog in *Homo sapiens*** |
| --- | --- | --- | --- | --- | --- | --- | --- | --- |
| Dynein light chain 4 | TTHERM_00716250 | I7MCM8 | 12.080 | 3.057 | 0.001 | 0.300 | NA | NA |
| Cell surface immobilization antigen SerH6, putative | TTHERM_00491130 | Q23J59 | 14.630 | 3.516 | 0.001 | 0.200 | [A0A250XQN9](https://www.uniprot.org/uniprot/A0A250XQN9) | NA |
| CAMP-dependent kinase, regulatory protein | TTHERM_00623090 | Q240X5 | 31.120 | 6.484 | 0.003 | 0.200 | NA | NA |
| Tetratricopeptide repeat protein | TTHERM_00313720 | Q22KA9 | 7.900 | 1.592 | 0.006 | 0.200 | IFT70 | [Q8N4P2](https://www.uniprot.org/uniprot/Q8N4P2) |
| CGMP-dependent kinase 5-1 | TTHERM_00046530 | Q23DN8 | 1.860 | 0.357 | 0.025 | 0.200 | NA | NA |
| Glutathione S-transferase, amine-terminal domain protein | TTHERM_00602870 | Q22YK8 | 6.730 | 1.172 | 0.000 | 0.200 | NA | [P09488](https://www.uniprot.org/uniprot/P09488) |
| Heat shock 70 kDa protein | TTHERM_00105110 | Q234I5 | 5.110 | 2.042 | 0.001 | 0.200 | NA | NA |
| Uncharacterized protein | TTHERM_00455660 | I7MI64 | 2.320 | 0.403 | 0.035 | 0.200 | NA | NA |

**Supplementary Table 7.** Proteins downregulated at least twofold in the *MEC17-KO* mutant compared to the *WT.*

| **Gene name** | **Gene ID** | **UniProt ID** | **Avg WT** | **Avg *MEC17-KO*** | **p value** | **Average fold change** | **Homolog in *C. reinhardtii*** | **Homolog in *Homo sapiens*** |
| --- | --- | --- | --- | --- | --- | --- | --- | --- |
| CAMP-dependent kinase catalytic subunit | TTHERM_00433420 | Q231B5 | 14.860 | 3.669 | 0.000 | 0.200 | NA | NA |
| CAMP-dependent kinase, regulatory protein | TTHERM_00623090 | Q240X5 | 31.120 | 7.513 | 0.003 | 0.200 | NA | NA |
| Tetratricopeptide repeat | TTHERM_00313720 | Q22KA9 | 7.900 | 1.471 | 0.000 | 0.200 | A8ITN7 | Q8N4P2 |
| Uncharacterized protein | TTHERM_00455660 | I7MI64 | 2.320 | 0.359 | 0.029 | 0.200 | NA | NA |
| Metallo-beta-lactamase family protein | TTHERM_00128670 | I7M1D2 | 9.530 | 1.418 | 0.004 | 0.100 | NA | NA |
| Intraflagellar transporter-like protein | TTHERM_00648910 | I7LT74 | 3.250 | 0.359 | 0.015 | 0.100 | IFT54 | IFT54 |
| Uncharacterized protein | TTHERM_01106190 | Q22BD8 | 4.410 | 0.359 | 0.011 | 0.080 | NA | NA |
| Intraflagellar transporter-like protein, putative | TTHERM_00149230 | I7LWB4 | 5.800 | 0.412 | 0.005 | 0.070 | IFT74 | IFT74 |
